# Exploring the diversity and community structure of the Testudines fecal mycobiome

**DOI:** 10.64898/2026.04.22.720109

**Authors:** Taylor Mills, Julia M. Vinzelj, Emma R. Cook, Emma Mills, Alexander J. Rurik, Jason W. Dallas, Donald M. Walker, Paul A. Stone, Cameron D. Siler, Mostafa S. Elshahed, Noha H. Youssef

## Abstract

Most gut microbiome studies have focused on bacteria, leaving a knowledge gap regarding gut associated fungi. We assessed fungal diversity in the gastrointestinal tract of the reptilian order Testudines (turtles and tortoises) using samples from 6 families, 19 genera, and 27 species. A highly diverse community affiliated with 17 phyla and 157 orders was encountered, with four phyla (Neocallimastigomycota, Chytridiomycota, Ascomycota, and Basidiomycota) representing 89.13% of the community. Neocallimastigomycota was identified in host families *Testudinidae* (land tortoises), *Chelidae*, *Chelydridae*, *Emydidae*, *Geoemydidae*, and *Kinosternidae*, with higher relative abundances in *Testudinidae* (40.18±37.97%) compared to all other families combined (2.71±4.04%). Neocallimastigomycota sequences were mostly affiliated with orders Testudinimycetales in the host family *Testudinidae* and Neocallimastigales in other host families. Chytridiomycota was identified in all host families, but was more ubiquitous and abundant in *Kinosternidiae* (45.17±34.12%), and exhibited a high level of variability across samples. Dikarya communities were highly diverse, with 108 orders identified, and, similar to Chytridiomoycota, exhibited a highly stochastic distribution pattern. Representatives of multiple yet-uncultured phyla (*Candidatus* “Algovoracomycota”, “Sedimentomastigomycota”, “Tartumycota” and “Cantoromastigomycota”) were identified, as well as eight novel orders in Chytridiomycota and Rozellomycota. Deterministic selection shaped community assembly in the host family *Testudinidae*, while the process was more stochastic in other host families. Distinct community structure was driven by differences in abundance and identity of the Neocallimastigomycota when comparing *Testudinidae* to. Our results describe a diverse and dynamic fungal community, shaped by the co-occurrence of autochthonous (resident) and transient (allochthonous) members of the gut microbiome.

**Importance:** Fungi are known to inhabit the gastrointestinal tract (GIT) of humans and mammals. However, information on the fungal community in the GIT of reptiles is relatively sparse. We investigated the diversity and community structure of fungi in the reptilian order Testudines. We conducted a culture-independent diversity survey on fecal samples obtained from 27 different host species. We identify representatives of 17 fungal phyla. As well, we demonstrate that the anaerobic gut fungi (phylum Neocallimastigomycota) are not restricted to the family *Testudinidae* (land tortoises) as previously suggested, but could successfully colonize and inhabit all other testudines families, including those exhibiting a predominantly omnivorous or carnivorous lifestyles. In addition, we expand on the known fungal diversity by identifying additional representatives of multiple recently described yet-uncultured phyla, and describe multiple novel orders and classes within existing phyla. Collectively, this effort adds to the growing body of knowledge of mycobiomes in underexplored animal hosts.

## Introduction

The kingdom Fungi encompasses a bewilderingly diverse array of microscopic and macroscopic organisms, all of which are characterized by heterotrophic osmotrophy (1). Fungi engage in trophic relationships with plants, animals, and microbial partners (1–3). The vast majority of studies focused on fungal diversity have targeted free-living (mostly terrestrial, but some aquatic (4–8) and plant-associated (ectomycorrhizal, endomycorrhizal, rhizosphere, phyllosphere, and endophytic (9–12) ecosystems. In contrast, significantly fewer studies have investigated animal-associated fungal communities (13–16). Within animal mycobiome diversity studies, emphasis has traditionally been placed on model fungal–animal associations of biotechnological (17) or pathogenic (18–22) potential. Although there continues to be a paucity of information on animal-associated fungal diversity for a wide range of hosts, there exists a nascent body of literature on the topic, primarily in mammals (14, 23–28). As such, a detailed understanding of the diversity and community structure of fungal communities in non-mammalian hosts is currently lacking.

Reptiles (Class Reptilia) represent a paraphyletic class in the Kingdom Animalia comprising the Squamata (lizards, snakes and amphisbaenians), Rhynchocephalia (tuatara), Testudines (turtles and tortoises), and the Crococdilians (29, 30). Despite recent efforts (31), the fungal community in reptiles remains poorly understood. This can be attributed to a number of factors, including their large global diversity (nearly 12,000 species, (32)), the restricted geographical distribution of many species (33, 34), and the propensity of some to be elusive or take refuge (35–37), impeding sample collection.

Within reptiles, the monophyletic order Testudines (comprising turtles and tortoises) is characterized by possessing bony shells developed from their ribs (38, 39). A wide range of habitat preferences, diet, and digestive strategies exist among extant Testudines. Ancestors of stem turtles are believed to have been terrestrial with some explorations of the aquatic medium (40–42). Currently, only tortoises (family *Testudinidae*) are purely terrestrial throughout their lives, with some members of the families *Emydidae* and *Geoemydidae* being semiaquatic/ terrestrial. Regarding diet preferences, members of family *Testudinidae* are herbivores throughout their adult lives. On the other hand, several freshwater and semiterrestrial turtles undergo an ontogenetic shift in their diet, where juveniles consume a mainly carnivorous (insectivorous) diet with high protein content for rapid growth requirements, becoming omnivores as they age (43, 44). A few species switch to a completely herbivorous diet, including the marine green sea turtle and a few species from the family *Emydidae* (cooters and sliders) (45–47). A notable exception is the *Chelydridae* (snapping turtles) and most side-necked turtles (e.g. *Pelusios* spp.), which do not undergo an ontogenetic shift, and remain carnivorous throughout their lives (43). While all Testudines possess a monogastric digestive system, with the large intestine being the site of microbial fermentation of carbohydrates to short chain fatty acids (48–51) in both omnivorous and herbivorous turtles (with few exceptions (52)), differences exist among host families with respect to food retention time, large intestine length, and the presence of enlarged dedicated fermentation chambers (e.g. enlarged caeca). These features increase with the level of herbivory in hosts (53–61).

Several studies have attempted to characterize the diversity and community structure of the Testudines gut microbiome and its role in the broader digestive processes. However, most Testudines gut microbiome studies have focused on bacteria (62–66), and neglected the gut fungi, despite their known importance in microbiome function (67–69). Recently, we conducted a targeted investigation on the occurrence and diversity of anaerobic gut fungi, phylum Neocallimastigomycota, in the herbivorous tortoises (family *Testudinidae*) (26). The identification of novel, tortoise-affiliated Neocallimastigomycota taxa in this study, and subsequent isolation of their representatives (70, 71) demonstrate that Testudines potentially harbor distinct fungal communities. Describing the Testudines gut mycobiome will broaden our understanding of the evolutionary history of Testudines–fungal trophic relationships, the role of sequestration in animal hosts in shaping fungal evolution, and the dynamics of fungal community acquisition and assembly in Testudines.

In this study, we examine the fungal community in 71 fecal/cloacal samples representing 6 different Testudines families to: 1) characterize the diversity and community structure of the Testudines mycobiome, 2) assess new fungal biodiversity at higher taxonomic ranks, and 3) identify factors impacting fungal community structure, with special emphasis on understanding the potential origin (resident versus transient) of various members of the community.

## Materials and Methods

### Ethics statement

Fecal samples from 70 Testudines were obtained between May 2023 and March 2025 post deposition by the animal. Zoo samples were obtained by trained personnel as part of the routine process of cleaning their habitat from deposited feces. Opportunistic sampling from ponds around OK, NM, TN, and LA were obtained with appropriate permits (MTSU IACUC permit no. 22-3002, TDEC permit no. 2025-083, TWRA permit no. 5107, ODWC permit no. 436044, NMDGF permit no. 2905). Additionally, a single cloacal swab sample was from a turtle in Camarines Norte province in Luzon, Philippines in strict accordance with the regulations established by the University of Oklahoma’s Institutional Animal Care and Use Committee (IACUC permit no. R17-019; R20-015; R21-005; and 2024-0272).

### Samples

Seventy-one samples were included in the analysis (Table S1). Included samples comprise representatives of both subgroups of Testudines; four Cryptodira (hidden-necked turtles) families (*Testudinidae* (n=42), *Emydidae* (n=12), *Kinosternidae* (n=10), *Geoemydidae* (n=4), and *Chelydridae* (n=2)), and one Pleurodira (side-necked turtles) family (*Chelidae* (n=1)), collectively belonging to 9 genera, and 27 species (Table S1). All *Testudinidae* (tortoise) samples were obtained from zoos (with the exception of 1 house pet). On the other hand, only 7 of the 29 other turtle samples originated from zoos with the remaining samples originating from opportunistic sampling on the landscape (Table S1). Testudines included in this study exhibit a wide range of feeding preferences (Table S1) including: 1. Preferentially carnivorous (plant matter constitutes <30% of diet and including plant matter only when the availability of animal food is limited), such as members of the family *Kinosternidae* (Sonora mud turtle and common musk turtle). 2. Omnivores with a carnivore preference (plant matter constitutes 30–50% of diet), such as members of the family *Chelydridae*. 3. True omnivores (plant matter constitutes 50–60% of diet), such as some members of the families *Emydidae* and *Geoemydidae*. 4. Omnivores with an herbivore preference (plant matter constitutes 60–75% of diet), including red-eared sliders and some members of *Testudinidae*. 5. Primarily herbivores (plant matter constitutes 75–90% of diet), including the Aldabra giant tortoise, red-footed tortoise, Texas tortoise, Asia brown tortoise, flat-backed spider tortoise, ploughshare tortoise. 6. Strict herbivores (plant matter constitutes >90% of diet), including the radiated tortoise, African spurred tortoise, giant tortoise, leopard tortoise, Russian tortoise, and Egyptian tortoise.

### DNA extraction, PCR amplification, and sequencing

DNA extraction from fecal samples was conducted either using DNeasy Plant Pro Kit (Qiagen Corp., Germantown, Maryland, USA) according to the manufacturer’s instructions as previously described (24, 26), or DNeasy 96 PowerSoil Pro (Qiagen Corp) following the manufacturer’s protocol. For PCR amplification, primers targeting the D2 region of the LSU rRNA in fungi were used LF402Fmix3 (5’-GTGAAATTGTCAAAAGGGAA-3’) and LR3 (5’-CCGTGTTTCAAGACGGG-3’). This primer pair was recently shown to effectively recover both Dikarya and non-Dikarya fungal lineages, compared to an observed bias towards Dikarya associated with primers targeting the ITS region (72). The primer pair targets a ∼350 bp region of the LSU rRNA gene and was modified to include the Illumina overhang adapters. PCR reactions used DreamTaq 2X Master Mix (Life Technologies, Carlsbad, California, USA) according to manufacturer’s instructions and contained 2 µl of DNA, and 2 µl of each primer (10 µM) in a 50 µl reaction mix. The PCR protocol included an initial denaturation for 5 min at 95 °C followed by 40 cycles of denaturation at 95 °C for 1 min, annealing at 55 °C for 1 min and elongation at 72 °C for 1 min, and a final extension of 72 °C for 10 min. PCR products were individually cleaned to remove unannealed primers using PureLink PCR cleanup kit (Life Technologies) according to manufacturer’s instructions. Cleaned products were then subjected to a barcoding second reaction to attach the Illumina UD indices (Illumina Inc., San Diego, California, USA) according to manufacturer’s instructions. These indexed PCR products were then cleaned to remove primers as well as any non-specific PCR products shorter than 200 bp using KAPA Pure Beads (Kapa Biosystems).

Cleaned products were individually quantified and pooled using the Illumina library pooling calculator (https://support.illumina.com/help/pooling-calculator/pooling-calculator.htm) to prepare 10 nM libraries. Pooled libraries were sequenced at the One Health Innovation Foundation lab at Oklahoma State University (Stillwater, Oklahoma, USA) on a NextSeq 2000 using a 2×300 bp paired-end library.

### Sequence processing and database creation

Sequence data were processed following the Mothur MiSeq SOP (https://mothur.org/wiki/miseq_sop/) (73). Assembly of forward and reverse Illumina reads was conducted using *make.contigs* command. Following, low quality sequences (with ambiguous bases, homopolymers longer than 8 bp, or length shorter than 200 or longer than 370 bp) were removed using *screen.seqs*. Redundant sequences were then removed using *unique.seqs*.

For alignment and classification purposes, we downloaded the LSU database created by (72) that combined the RDP training set 11 and the Silva LSU v132 reference database (henceforth RDP+Silva). To overcome the rarity of reference sequences associated with the Neocallimastigomycota in this database, we supplemented the RDP+Silva database with sequences from the Anaerobic Fungi Network LSU_Database v2.0 (https://anaerobicfungi.org/databases/). The updated database (henceforth RDP+Silva_AGF) was used for classification using local Blastn (74) analysis as described below.

Non-redundant sequences were aligned to the RDP+Silva_AGF using *align.seqs* in Mothur. Misaligned sequences were removed, and *pre.cluster* was used to de-noise the sequences (at 1%). Chimeras were identified and removed (*chimera.vsearch*), and the remaining sequences were then clustered at 94% (genus equivalent). Singletons were removed and the remaining OTUs (n=2347) were then classified into genera as explained below.

### Phylogenetic analysis

OTU representatives were compared to the RDP+Silva_AGF database using local Blastn. We used a cutoff of 94% Blastn similarity to assign an OTU to a fungal genus. Sequences that remained unclassified (Blastn similarity to a top hit in the RDP+Silva_AGF database <94) were then compared to the NCBI nr (75) and the Eukaryome (76) databases using Blastn. Top hits (any hit with at least 90% similarity and at least 70% query coverage) were identified. Unclassified sequences and their top hits were then inserted into reference fungal trees with representatives of all fungal phyla. At this level, sequences that phylogenetically clustered within a known genus were assigned to that genus. Two exceptions from the above classifications were sequences that clustered into Chytridiomycota and Rozellomycota. We assigned Chytridiomycota sequences only at the order rank since genus-level delineation is not yet clear for the orders Lobulomycetales, Gromochytriales, Mesochytriales, Rhizophydiales, and Zygophlyctidales. Sequences that clustered within Chytridiomycota but not within an already described order were assigned to putative new order designations. For Rozellomycota-affiliated sequences, we assigned sequences to a class-level if they did not cluster into the described orders Paramicrospordiales, Morellosporales, or with the genus *Rozella*. Sequences that were assigned by Blastn against the Eukaryome database to any of the thirty new taxa proposed by (77) were given the order-level taxonomic designation of their first hit. However, since the International Code of Nomenclature for Algae, Fungi, and Plants (ICNafp) does not accept eDNA as the type material (78), we have added the notation “*Candidatus*” to the names proposed by (77) (Table S2). For these later sequences, our phylogenetic analysis mostly confirms the high-rank taxonomy presented in (77) with two exceptions; the position of *Candidatus* “Aquamastigomycetes”, *Candidatus* “Cantoromastigomycetes”, *Candidatus* “Dobrisimastigomycetes”, *Candidatus* “Palomastigomycetes”, and *Candidatus* “Sedimentomastigomycetes” as classes in the phylum Neocallimastigomycota, and the position of *Candidatus* “Algovoracomycetes” as a class in Monoblepharomycota. As (77) does not provide clear quantitative rank assignment thresholds; such assessment hence appears to be subjective and open to revision. In our analysis, reference sequences from the five proposed Neocallimastigomycota classes had variable positions in our trees, and mostly clustered as sister lineages to the Neocallimastigomycota and the Monoblepharomycota. Similarly, *Candidatus* “Algovoracomycetes” split from other members of Monoblepharomycota in our trees, as previously observed in (79). As such, we could not clearly confirm the affiliation of these classes with the Neocallimastigomycota, and the Monoblepharomycota, respectively. Further, all current members of the Neocallimastigomycota are strict anaerobes and are known to be exclusively host-associated (with herbivorous mammals, birds, or tortoises) (70, 71, 80–82). Sequences for the five newly proposed Neocallimastigomycota “classes” were obtained from aquatic and terrestrial environments that are aerobic in nature, which argues against their affiliation with the phylum. Similarly, members of the Monoblepharomycota are known to be saprobic on fruits and twigs in soil and aquatic ecosystems (78), while the *Candidatus* “Algovoracomycetes” (formerly recognized as NC_ChyL-1) members are believed to be epibionts on algae (79, 83). We, therefore, opted for assigning sequences that clustered with any of these six classes proposed by (77) to a phylum rather than a class rank (Table S2).

### Statistical analysis and correlation to host traits

We used the non-parametric Kruskal-Wallis test followed by post-hoc Wilcoxon with Bonferroni-Hochberg correction for pairwise comparisons to identify host factors predictive of the four major fungal phyla (Neocallimastigomycota, Chytridiomycota, Ascomycota, and Basidiomycota). Effect sizes for Kruskal-Wallis tests were calculated based on the H-statistic obtained and multiplied by 100.

Five host factors were considered and are all shown in Table S1: host family, host food retention time (short, ∼10–12 days; medium, 15–20 days; and long, >20 days), feeding preferences as explained above, host adult carapace length (small, <20 cm; medium, 20–80 cm; and large, >80 cm), and domestication status.

For Neocallimastigomycota community, we used differential abundance analysis with bias correction (using ANCOM-BC2) (84, 85) on genus-level counts. We used a one-versus-all design to compare each host family to all other samples. Bias-corrected log fold changes were estimated for each genus, and statistical significance was determined using Benjamini–Hochberg adjusted p-values (q < 0.05). For host families with detected differentially abundant taxa, volcano plots were constructed in R using ggplot2 (86).

### Role of stochastic versus deterministic processes in fungal community assembly

We used two approaches (the normalized stochasticity ratio (NST) (87), and the null-model-based quantitative framework implemented by (88, 89)) to examine the contribution of various deterministic (niche theory-based) and stochastic (null theory-based) processes (87–89) in shaping the fungal community assembly. Due to relatively low number of replicates from some hosts, we restricted our analysis to three host families with n ≥ 10 samples (*Testudinidae*, *Kinosternidae*, and *Emydidae*). As well, we compared six different host species with n ≥ 5 individuals (*Aldabrachelys*, *Astrochelys*, *Centrochelys*, *Chelonoidis*, *Kinosternon*, and *Trachemys*).

For calculating NST, we used the NST package in R (Version 3.1.10; (87)). NST was calculated based on both the incidence-based Jaccard index and the abundance-based Bray-Curtis index, where an NST value of >50% indicates a more stochastic assembly, while values <50% indicate a more deterministic assembly. Following, to test the significance of difference between pairs, we used *<u>nst.boot</u>* in the R NST package to randomly draw samples within each comparison group, followed by bootstrapping (n=1000) of NST values. Values obtained after bootstrapping were then compared using Wilcoxon test with Benjamini-Hochberg adjustment.

To further quantify the contribution of specific deterministic (homogenous and heterogenous selection) and stochastic (homogenizing dispersal, dispersal limitation, and drift) processes in shaping the fungal community assembly, the iCAMP R package (Version 1.5.12; (90)) was used to calculate beta net relatedness index (βNRI) and modified Raup-Crick metric based on Bray-Curtis metric (RC_Bray_) using the function *<u>bNRIn.p</u>*. Following, the values of βNRI and RC_Bray_ were used to partition selective processes into homogenous and heterogenous selection and stochastic processes into homogenizing dispersal, dispersal limitation, and drift, as detailed before in (24), and the percentages of pairwise comparisons falling into each category were used as a proxy for the contribution of each of these processes to the total fungal community assembly.

### Fungal community structure in Testudines

Weighted Unifrac was calculated for all possible pairs of samples using the function *ordinate* in the Phyloseq R package (Version 1.50.0; (91)), and was used to construct principal coordinate analysis plots using the function *plot_ordination*. We used PERMANOVA tests (run using the function *adonis2* in the Vegan R package (Version 2.7–2; (92)) to partition the dissimilarity among the sources of variation (including animal host species, animal host family, animal food retention time, diet description, carapace size, and domestication status), and the F-statistic, p-values, and R^2^ values were compared to identify the factors that significantly affect the AGF community structure. We used interaction terms in the model to account for interdependence between host factors (e.g. feeding preference and retention time, host genus and feeding preference, and host family and domestication since almost all *Testudinidae* samples were obtained from zoo settings, while other families were mainly sampled from the wild). The percentage of the sum of squares of each factor to the total sum of squares was used to calculate the percentage variance explained by each factor. Additionally, we used the function ordinate in the Phyloseq R package to calculate Bray Curtis beta diversity indices for all possible pairs of samples, followed by constructing a double principal coordinate analysis (DPCoA) plot (using the function plot_ordination), which allows both the samples and the taxa to be plotted on the same coordinate space. Taxa with Euclidean distances close to group centroids are considered to contribute more to the community structure of the group.

### GenBank Accession

Illumina reads were deposited in NCBI SRA under BioProject accession number PRJNA1422821.

## Results

### Fungal community overview

The fungal community observed in Testudines was highly diverse, with representatives of 17 phyla and 157 orders identified (Figure 1, Table S2). Broadly, the community was dominated by sequences affiliated with the anaerobic gut fungi (phylum Neocallimastigomycota), Chytrids (phylum Chytridiomycota), and Dikarya (phyla Ascomycota and Basidiomycota). These four phyla, collectively, represented 89.13% of all sequences (Figure 1A) and an average of 93.14±16.59% (average ± standard deviation) in individual samples (Table S2). Additional non-Dikarya phyla identified were present as minor components in most samples (Figure 1, Table S2). Many non-Dikarya phyla encountered have cultured representatives (Mucoromycota, Monoblepharomycota, Basidiobolomycota, Blastocladiomycota, Aphelidiomycota, Rozellomycota, Mortierellomycota, Olpidiomycota, Entomophthoromycota, and Glomeromycota), but some are phyla with yet-uncultured representatives that have been named using eDNA as the type material, e.g., *Candidatus* “Algovoracomycota”, *Candidatus* “Cantoromastigomycota”, *Candidatus* “Sedimentomastigomycota”, and *Candidatus* “Tartumycota” (77) (Figure 2).

**Figure 1.**
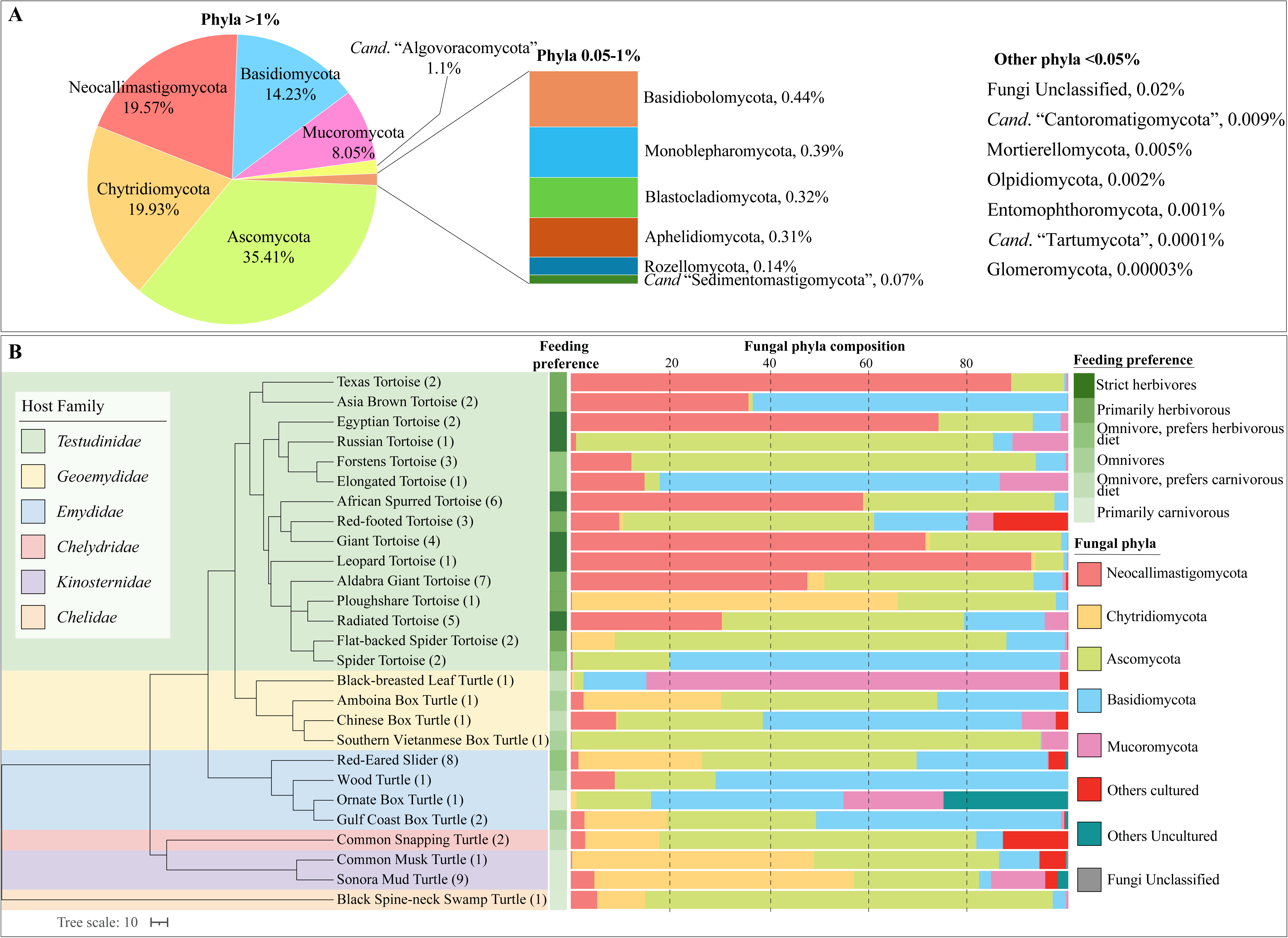
Overview of the fungal community in the Testudines gut. (A) Phylum-level composition of all sequences encountered in the 71 samples studied. (B) Phylum-level composition per host species. The phylogenetic tree showing the relationship between host species was downloaded from timetree.org, and the number of individuals belonging to each species is shown in parentheses following the species name. The track to the right of the tree shows the host feeding preferences. Fungal community composition for each animal species is shown to the right as colored columns. “Others cultured” include Monoblepharomycota, Basidiobolomycota, Blastocladiomycota, Aphelidiomycota, Rozellomycota, Mortierellomycota, Olpidiomycota, Entomophthoromycota, and Glomeromycota, while “Others uncultured” include *Candidatus* “Algovoracomycota”, *Candidatus* “Cantoromastigomycota”, *Candidatus* “Sedimentomastigomycota”, and *Candidatus* “Tartumycota”. Detailed fungal composition of every sample is presented in Table S2.

**Figure 2.**
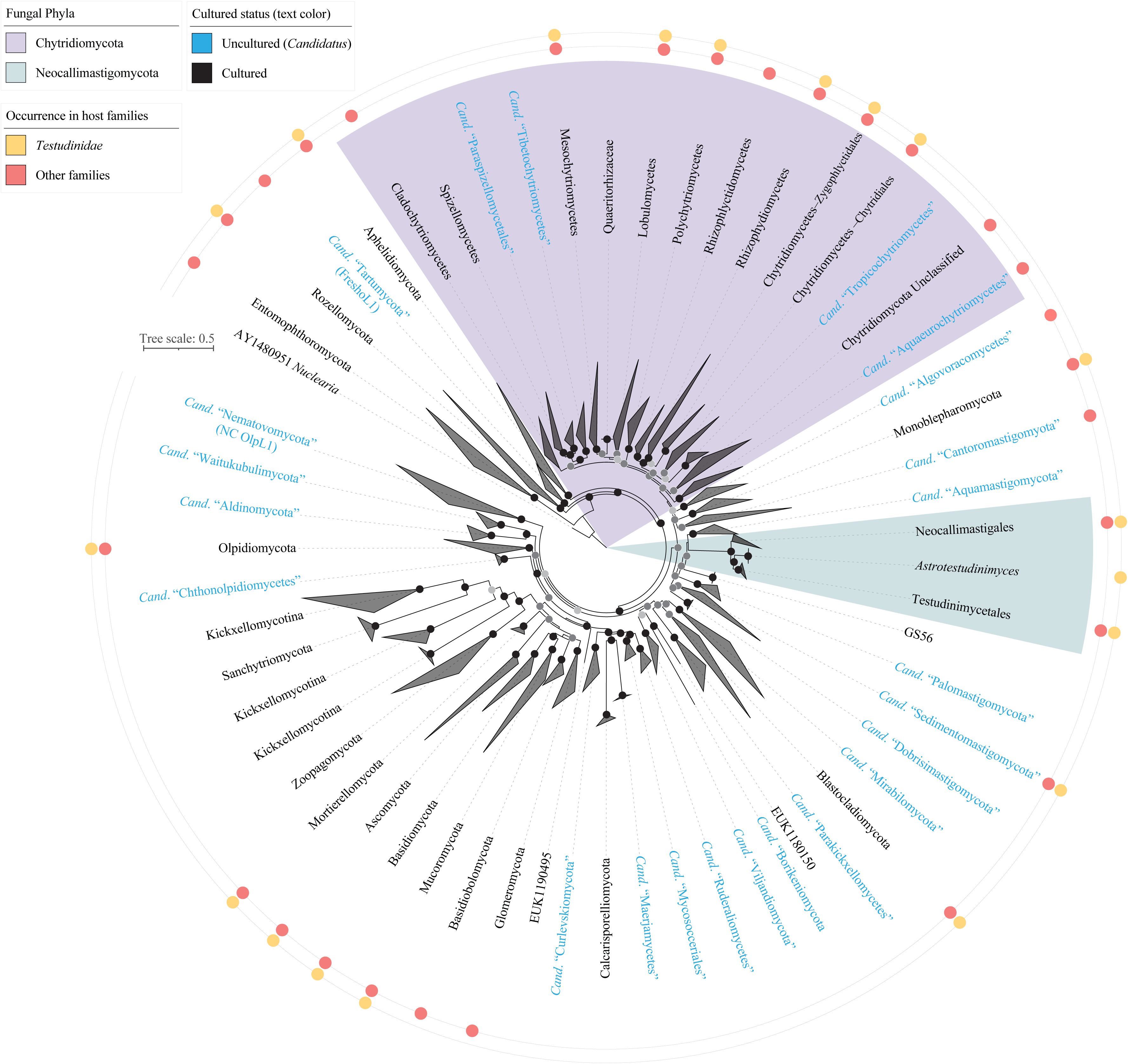
Maximum likelihood phylogenetic fungal tree of life based on the alignment of the D1–D2 regions of the LSU rRNA. The tree includes representatives from all previously reported cultured, and uncultured phyla as references. Recently proposed uncultured taxa in (77) are shown in cyan text and are designated as *Candidatus*. The two tracks around the tree indicate whether representatives of the phyla were encountered in the host family *Testudinidae* (land tortoises) (yellow circles), or other turtle families (salmon circles). The two orders in Neocallimastigomycota and *Astrotestudinimyces* genus Incerta sedis are shown. Similarly, class-level is shown for Chytridiomycota. The three bootstrap support values (SH-aLRT, aBayes, and UFB) are shown as colored dots as follows: all three support values > 70%, black dot; 2/3 support values > 70%, dark grey; 1/3 support values > 70%, light grey.

### Neocallimastigomycota community in Testudines

Sequences affiliated with the *Neocallimastigomycota* represented 19.56% of the entire dataset (Figure 1A), and were identified in nearly all (68/71) samples examined. Neocallimastigomycota relative abundance ranged between 0.01-99.85% (Figures 1B, 3, Table S2), and was higher in family *Testudinidae* (tortoises) (42.36±37.99%), compared to all other host families (3.31±4.28%) (Student T-test p-value =9.7×10^-7^). Furthermore, Neocallimastigomycota represented >10% of the total sequences in 27/42 of family *Testudinidae* samples, compared to only in 2/29 of all other host families (Figures 1, 3, Table S2). The relative abundance of Neocallimastigomycota was significantly (Kruskal Wallis p <0.002) and positively associated with feeding preferences (Kruskal effect size measure = 31.9%), food gut retention time (29.7%), host family (16.9%), carapace length (15.5%), and domestication status (14.6%) (Table S3, Figure S1A). Pairwise comparisons of different feeding preferences showed species with a primarily herbivorous diet harbored significantly more Neocallimastigomycota than those that are primarily carnivorous or omnivorous with a carnivore diet preference (Wilcoxon test p-value <0.05), while strict herbivores had significantly more Neocallimastigomycota than all other feeding preferences (except primarily herbivores) (Wilcoxon test p-value <0.05). Similarly, pairwise comparisons of Neocallimastigomycota relative abundances showed a significantly higher abundance in *Testudinidae* compared to *Emydidae* (p-value = 0.03), in Testudines with long food retention time (>20 days, p-value <0.034), in Testudines with larger carapace size (>80 cm, p-value <0.04), and in Testudines from Zoo settings (p-value = 0.001) (Table S3).

**Figure 3.**
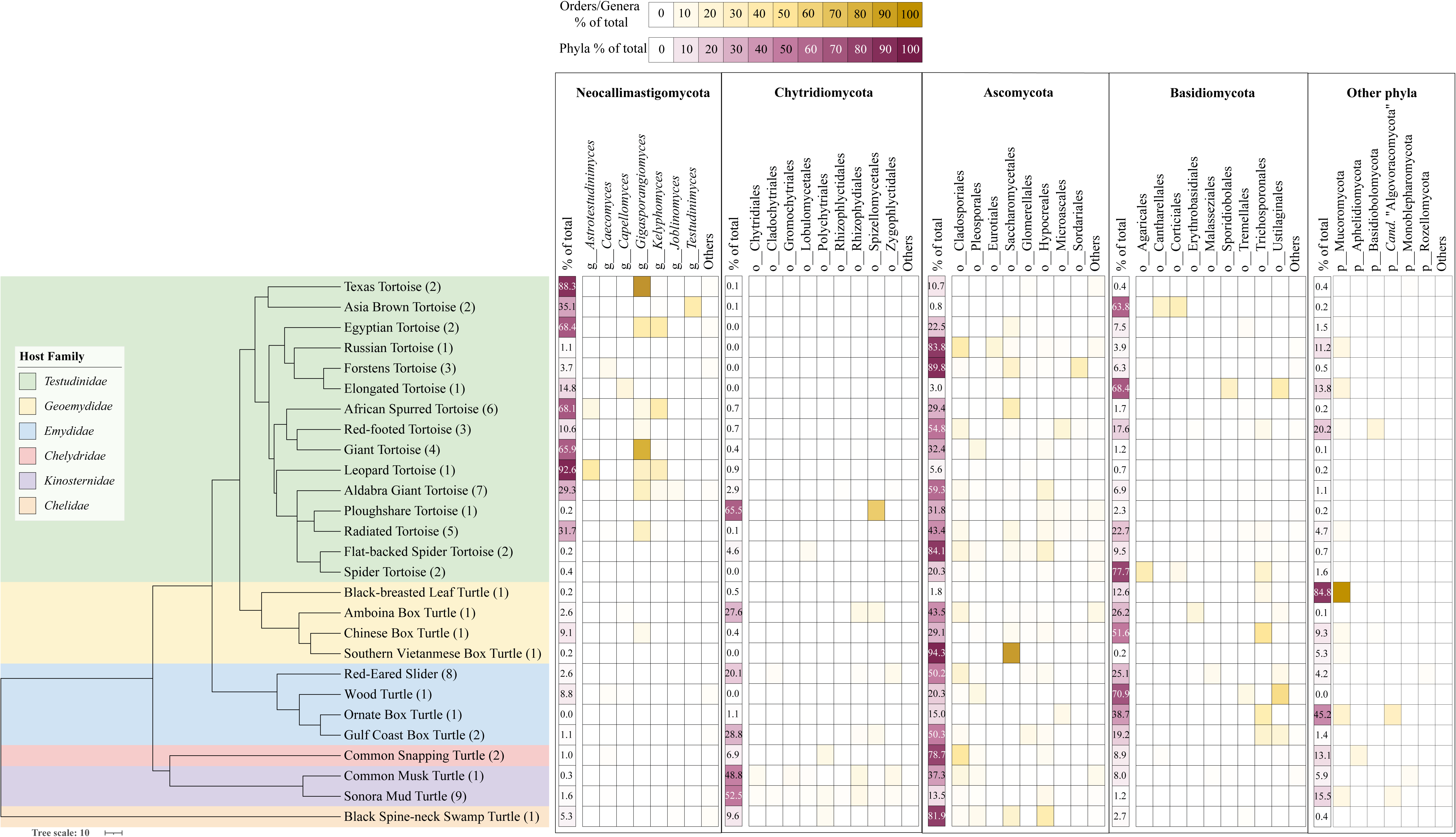
Lower-taxa composition of the four most abundant fungal phyla. The phylogenetic tree showing the relationship between turtle species was downloaded from timetree.org, and the number of individuals belonging to each animal species is shown in parentheses following the species name on the branches. For each of the four phyla, first a heatmap (shades of pink) is shown depicting the number of sequences belonging to that phylum as a percentage of the total sequences (header % of total) followed by another heatmap (shades of yellow) depicting the lower-taxa composition distribution. Lower taxa are shown at the genus level for Neocallimastigomycota, and at the order level for Chytridiomycota, Ascomycota, and Basidiomycota. Numbers are averages from all individuals belonging to each host species. Only taxa whose composition exceeded 10% on average in at least one host species are shown. All other taxa are grouped under “Others”. To the very right, heatmaps are shown for phyla other than the major four. For these phyla, heatmaps are based on phylum level composition. Only phyla whose composition exceeded 10% on average in at least one host species are shown. All other phyla are grouped under “Others”.

In addition to differences in relative abundance, a distinct community composition pattern was observed. In the Family *Testudinidae* (tortoises), >50% of the Neocallimastigomycota community in the majority of samples (31/42) was represented by genera *Gigasporangiomyces*, *Kelyphomyces*, *Astrotestudinimyces*, and *Testudinimyces* in the order *Testudinimycetales* (recently proposed to accommodate taxa isolated from tortoises (70, 71)) (Figure 3, Tables S2, S4). On the other hand, the Neocallimastigomycota community in samples from all other host families (26/29) was predominantly affiliated with genera in the *Neocallimastigales*, the order that comprises all genera previously isolated from mammalian sources (81). These include the genera *Caecomyces*, *Capellomyces*, *Neocallimastix*, and, notably, *Joblinomyces*, a genus with relatively rare distribution and consistently comprising an extremely minor fraction of the Neocallimastigomycota community in mammalian fecal samples examined (24, 25, 93). Analysis of the Neocallimastigomycota community composition using ANCOM-BC2 confirmed the significant enrichment of the Testudinimycetales genera *Gigasporangiomyces*, *Kelyphomyces*, and *Testudinimyces* in members of Family *Testudinidae* (tortoises), with the Neocallimastigales genera *Neocallimastix*, *Khoyollomyces*, *Anaeromyces*, *Orpinomyces*, and *Caecomyces* significantly enriched in other host families (Figure S2A).

Limited novelty was observed in the Neocallimastigomycota, with sequences potentially representing one novel genus in the order Neocallimastigales (0.09±0.17% in 4 samples), and another novel genus in the order Testudinimycetales (0.002±0.001% in 15 samples) both identified as minor fractions in this study (Figure S3).

### Chytridiomycota community in Testudines

Sequences associated with the phylum Chytridiomycota represented 19.93% of the entire dataset, and were identified in 49/71 samples examined, with individual relative abundance ranging between 0.0004–92.89% (18.68±26.6%) (Figures 1, 3, Table S2). In contrast to the Neocallimastigomycota, the relative abundance of the Chytridiomycota was significantly lower in Family *Testudinidae* (tortoises) (occurrence, 23/42 samples; abundance, 5.09±14.1%) (Student t-test p-value = 0.0005) compared to all other host families (occurrence, 27/29 samples; abundance, 30.27±29.36%). The family *Kinosternidae* (occurrence, 10/10 samples; relative abundance 51.43±30.55%) harbored more Chytridiomycota, followed by the family *Emydidae* (occurrence, 11/12 samples; relative abundance 21.37±23.84%), with the remaining three families harboring significantly lower Chytridiomycota relative abundance (occurrence 6/7 samples; average relative abundance 11.3±12.72%) (Student test p-value<0.02).

Out of the five host factors tested, food retention time (Kruskal effect size measure = 19.4%), host family (16.6%), and domestication status (14.6%) significantly correlated with Chytridiomycota relative abundance (Kruskal Wallis p-value <0.01). Pairwise comparison showed that Testudines with short food retention time (<12 days) harbored significantly more Chytridiomycota (p-value < 0.005), *Emydidae* harbored significantly higher Chytridiomycota than *Testudinidae* (p-value = 0.01), and wild Testudines harbored significantly more Chytridiomycota than zoo-housed samples (p-value =0.0009) (Table S3, Figure S1B).

While representatives of 15 Chytridiomycota orders were identified in the samples, only 9 were dominant (>10% relative abundance in at least 1 sample) (Figure 3). Within the family *Kinosternidae*, sequences affiliated with order *Rhizophydiales* dominated in replicates of *Kinosternon sonoriense* (Sonora mud turtles) followed by sequences affiliated with *Spizellomycetales* and *Chytridiales*, while the single *Sternotherus odoratus* (Common musk turtle) sample harbored mainly *Rhizophydiales* and *Zygophlyctidales* sequences. *Rhizophydiales*, *Spizellomycetales*, and *Zygophlyctidales* were also abundant in the family *Emydidae* especially *Terrapene carolina* (common box turtle) and *Trachemys scripta* (red-eared slider). In *Testudinidae* (tortoises), Chytridiomycota sequences were less abundant with only three samples harboring a Chytridiomycota order with relative abundance >10% (*Spizellomycetales* in *Astrochelys yniphora* (ploughshare tortoise), *Zygophlyctidales* in *Aldabrachelys gigantea* (Aldabra giant tortoise), and *Lobulomycetales* in *Pyxis arachnoides* (spider tortoise)).

While the majority of Chytridiomycota sequences identified were associated with known orders with cultured representatives, a minor fraction was affiliated with novel, yet-uncultured lineages. Specifically, we identified two new orders in class Mesochytriomycetes (Mesochytriomycetes_Ord01, Mesochytriomycetes_Ord02) (Figure S4). These orders have close representatives in soil and aquatic sediment, and were labeled as Mesochytriomycetes.ord01 and Mesochytriomycetes.ord02 in the Eukaryome database (76). While representing an overall minor fraction of all Chytridiomycota sequences, Mesochytriomycetes_Ord01 was present in relatively high abundances in two *Kinosternon sonoriense* (Sonora mud turtles) (abundances 17.31% and 1.24%).

### Ascomycota community in Testudines

Ascomycota was the most abundant phylum, constituting 35.41% of the entire dataset (Figure 1A), and a significant fraction (>10% of sequences) in 49/71 samples. Ascomycota relative abundance was similar across host families and showed no correlation to feeding preference, carapace size, feed retention time, or domestication status (Table S3). Given the immense diversity of Ascomycota in nature, it is not surprising that representatives of 78 different orders were identified in the dataset (Table S2). However, only 8 orders were dominant (>10%) in one or more samples (*Cladosporiales*, *Pleosporales*, *Eurotiales*, *Saccharomycetales*, *Glomerellales*, *Hypocreales*, *Microascales*, and *Sordariales*) (Figure 3).

These orders collectively constituted >70% of all Ascomycota sequences in most samples (67/72). Interestingly, *Cuora picturata* (Southern Vietnamese box turtle) was dominated (98.46% of total sequences) with Saccharomycetales (single OTU, 99.22% sequence similarity to *Candida glabrata*. The four most abundant Ascomycota OTUs showed highest similarity to well-known genera (>99% similarity) (Table S4).

### Basidiomycota community in Testudines

Basidiomycota was the fourth most abundant phylum, constituting 14.22% of the entire dataset (Figure 1A). Basidiomycota sequences were encountered in 69 of the 71 samples and represented >10% in 27/69 samples. Host family significantly explained Basidiomycota relative abundances (Kruskal effect size measure 15.4%), with higher abundances encountered in *Emydidae* (34.88±28.17%) compared to *Testudinidae* (14.56±22.99%) (Student t-test p-value=0.013), and *Kinosternidae* (2.23±2.67%) (Student t-test p-value=0.0016), but not *Geoemydidae* (22.81±22.22%) (Student t-test p-value=0.45) (Figure S1C, Table S3).

Representatives of 47 Basidiomycota orders were identified in the samples studied (Table S2). Of these, nine were the most abundant (present in >10% of the dataset) including *Agaricales*, *Cantharellales*, *Corticiales*, *Erythrobasidiales*, *Malasseziales*, *Sporidiobolales*, *Tremellales*, *Trichosporonales*, and *Ustilaginales* (Figure 3). These orders collectively constituted >70% of all Basidiomycota sequences in the majority of samples (54/69). The four most abundant Basidiomycota OTUs showed highest similarity to well-known genera (>98% similarity) (Table S4).

### Additional phyla in Testudines

Representatives of 14 additional phyla were identified and collectively constituted 10.85% of the entire dataset (range 0.00003–8.05%, average 0.83±2.21%; Table S2; Figure 1). Of these, only six phyla were present in high relative abundance (>10%) in at least one sample. These include 1) Mucoromycota, specifically the genus *Mucor* that constituted up to 83.1% in one sample, 2) the newly proposed orders *Candidatus* Algovoracales” and *Candidatus* “Solivoracales” within the newly proposed phylum *Candidatus* “Algovoracomycota” (Figure S5) that constituted 24.94% and 70.11%, respectively in two samples, 3) Basidiobolomycota, specifically the genus *Basidiobolus* that constituted 44.32% in one sample, 4) Blastocladiomycota, constituting 10.65–11.73% in two samples, 5) Aphelidiomycota, specifically the family *Aphelidiaceae* constituting 9.45–25.18% in three samples, and 6) Rozellomycota, specifically the uncultured classes GS06, and GS137 that constituted 13.36%, and 20.29%, respectively, in two samples. Additional phyla were minor constituents (<0.1% of the total dataset) and included *Candidatus* “Sedimentomastigomycota”, *Candidatus* “Cantoromastigomycota”, Mortierellomycota, Olpidiomycota, Entomophthoromycota, *Candidatus* “Tartumycota”, and Glomeromycota (Table S2, Figure 1A).

### Stochastic and deterministic processes shaping the Testudines gut mycobiome

To identify the contribution of stochastic and deterministic processes to the fungal community assembly in Testudines, we first examined Normalized Stochasticity Ratios (NST) calculated based on two β-diversity indices (abundance-based Bray-Curtis index, and incidence-based Jaccard index) and compared them across the three host families with > 10 individuals (*Emydidae*, *Kinosternidae*, and *Testudinidae*), and genera with >5 individuals (*Trachemys*, *Kinosternon*, *Aldabrachelys*, *Astrochelys*, *Centrochelys*, and *Chelonoidis*). Regardless of the index used, NST values were significantly lower in family Testudinidae genera (Wilcoxon adjusted p-value <1.2×10^-6^ for all pairwise comparisons except *Kinosternon*–*Astrochelys*) (Table S5, Figure 4A–D), indicating a less stochastic community assembly in Testudinidsae (tortoises). Within the four tortoise genera, values of NST increased in the order: *Centrochelys*, *Aldabrachelys*, *Chelonoidis*, and *Astrochelys.* Additionally, the genus *Trachemys* showed significantly higher NST values compared to *Kinosternon*.

**Figure 4.**
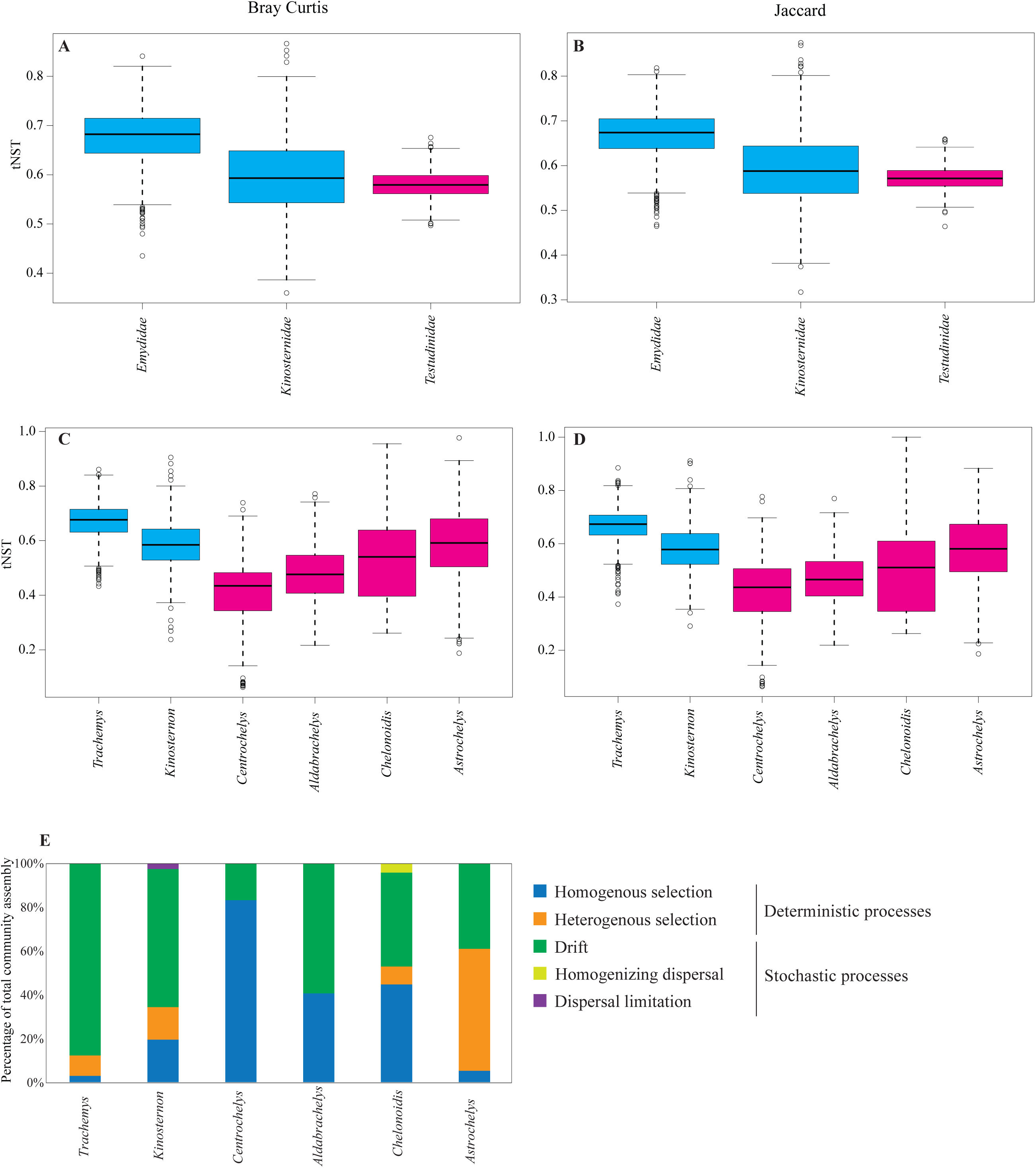
Contribution of stochastic and deterministic processes to the Testudines fungal community assembly. (A–D) Levels of stochasticity in the fungal community assembly were compared between different host families (A, B; for families with more than 10 individuals), and host genera (C, D; for genera with more than 5 individuals). Two normalized stochasticity ratios (NST) were calculated; the abundance-based Bray-Curtis index (A–C), and the incidence-based Jaccard index (B–D). (E) The percentages of the various deterministic and stochastic processes shaping the fungal community assembly in the six different animal species.

To quantify the contribution of specific deterministic (homogenous and heterogenous selection) and stochastic (homogenizing dispersal, dispersal limitation, and drift) processes in shaping the fungal community assembly in Testudines, we employed the previously suggested (87–89) two-step null-model-based quantitative framework. Results (Figure 4E) confirmed the more stochastic community assembly in *Emydidae*, *Kinosternidae* (stochasticity was 65.43% of the total community assembly in *Trachemys*, and 87.5% of the total community assembly in *Kinosternon*) as opposed to a more deterministic community in *Testudinidae* taxa (deterministic processes accounted for 40.82–83.33% of the total community assembly). More specifically, the fungal community in *Trachemys* and *Kinosternon* is shaped by drift. In *Testudinidae*, homogenous selection shapes the community in *Centrochelys*, *Aldabrachelys*, *Chelonoidis*, while heterogenous selection is responsible for community assembly in *Astrochelys* (Figure 4E).

### Gut fungal community structure in Testudines

We assessed the fungal community structure in the 71 samples studied using PCoA constructed based on the phylogenetic similarity-based beta diversity index weighted Unifrac (Figure 5A). PERMANOVA test showed that host genus, host family, food retention time, feeding preferences, carapace size, but not domestication status, significantly explained variance in fungal community structure (weighted Unifrac; F-statistic p-value <0.05), with host phylogeny and feeding preferences contributing to total variance more than other host factors (Figure 5A, Table S6). The interaction between feeding preference and food retention time was also found to be significant (F-statistic=2.449, p-value=0.035). The DPCoA ordination biplot (Bray Curtis index) (Figure 5B) showed separation of the tortoise family *Testudinidae* from the families *Emydidae*, *Kinosternidae*, and *Geoemydidae*. Such separation was mainly brought about by the prevalence of the Testudinimycetales genera *Gigasporangiomyces*, *Kelyphomyces*, and *Gopheromyces* and the genus *Astrotestudinimyces* in *Testudinidae* samples (Figure 5B, genera highlighted as grey squares). This effect was further apparent in DPCoA plots constructed using only Neocallimastigomycota OTUs (Figure S2B).

**Figure 5.**
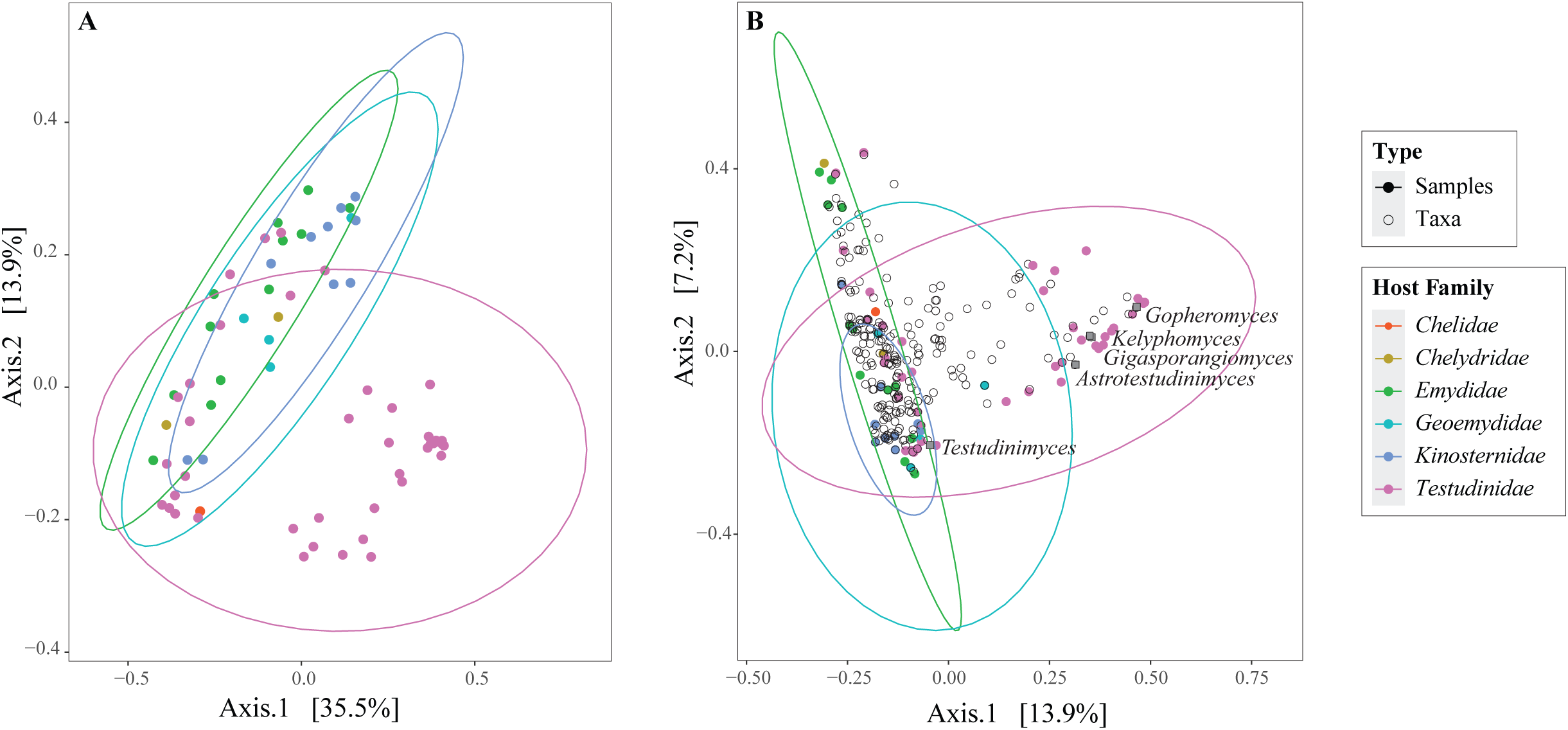
Gut fungal community structure in Testudines. (A) Principal coordinate analysis (PCoA) ordination plot based on the fungal community structure in the 71 samples studied here. PCoA was constructed using the phylogenetic similarity-based weighted Unifrac index. The percent variance explained by the first two axes is displayed on the axes. Samples are color-coded by animal family with ellipses encompassing 95% of variance for animal families with >4 individuals. (B) Double principal coordinate analysis (DPCoA) biplot based on the Bray Curtis index. The percentage variance explained by the first two axes is displayed on the axes. Samples are color-coded by animal family with ellipses encompassing 95% of variance for animal families with >4 individuals. Fungal genera are shown as black empty circles, and the four Testudinimycetales genera as well as the genus *Astrotestudinimyces* are labeled and shown as grey rectangles.

## Discussion

Our analysis of the Testudines gut fungal community adds to the growing body of knowledge of mycobiomes in underexplored animal hosts. Our study revealed high fungal diversity (17 phyla, 157 orders) in the Testudines GIT tract (Figure 3), with four phyla (Ascomycota, Chytridiomycota, Neocallimastigomycota, and Basidiomycota) representing the core community in most samples (Figure 1). In addition, we established the occurrence of the Neocallimastigomycota in all families of Testudines examined (Figures 1, 3). We also expanded on the fungal tree of life by identifying additional representatives of multiple recently described yet-uncultured phyla (*Candidatus* “Algovoracomycota”, *Candidatus* “Sedimentomastigomycota”, *Candidatus* “Cantoromastigomycota”, and *Candidatus* “Tartumycota”) (Figure 1), and identified novel orders, and classes in Chytridiomycota, and Rozellomycota, respectively (Figures S4, S6). We demonstrated that fungal assembly and community structure is deterministic in tortoises (family *Testudinidae*) compared to the primarily stochastic selection in families *Emydidae* and *Kinosternidae*) (Figure 4). We established animal host as the most important determinant for shaping Testudines microbial community structure, clearly separating family *Testudinidae* (tortoises) from other turtle families (Figure 5), and showed that such distinction in community structure is driven by the Neocallimastigomycota component of the community (Figure S2).

We argue that these patterns could best be understood considering differences in host microhabitat and feeding preferences, as well as by the physiological preferences and metabolic capacities of the gut mycobiome. In prior efforts, we identified (26) and described (70, 71) a novel community of Neocallimastigomycota in members of family *Testudinidae* (tortoises). The present study confirms the universal occurrence of Neocallimastigomycota in a wider range of *Testudinidae* hosts and, more importantly, identifies its presence in all five families and twelve species of Testudines examined, regardless of their feeding preference (Figure 1). The identification of Neocallimastigomycota in turtles challenges the prior assumption (based on studies in mammalian hosts) that Neocallimastigomycota is only capable of colonizing strict herbivores that possess a dedicated fermentation chamber (rumen, verticula, or enhanced caecum or colon) (80). While turtles in the family *Testudinidae* (tortoises) satisfy these two requirements, other turtle families are neither strictly herbivorous (Table S1), nor possess a dedicated fermentation chamber in their guts. Nevertheless, we note that all turtles exhibit a relatively long food residence time compared to mammals (at least three days (Table S1), compared to 20–50 h in mammalian ruminants (94) and 35–45 h in mammalian hindgut fermenters (95)), with anaerobic conditions prevailing in their large intestine. Turtles also include, to various degrees, a plant component in their diet (Table S1). We thus argue that long feed retention time, anaerobic conditions, and a plant component in the diet enable Neocallimastigomycota colonization of resident plant material in the GIT of turtles. The relative success in colonization and propagation of Neocallimastigomycota in turtles could depend on the prevailing temperature, since turtles are poikilothermic. Members of the order Neocallimastigales identified in this study are known to have a relatively narrow temperature growth range (37–42°C; (81)). These temperatures are generally at the upper end of the thermal breadth for many turtle species (96–99), suggesting the potential for unique thermal adaptations of Neocallimastigales to persist in turtles.

Nevertheless, the distinct differences in physiological and metabolic properties between members of the order Testudinimycetales and genus *Astrotestudinimyces* (prevalent in family *Testudinidae*) and Neocallimastigales (prevalent in other turtle families) hint to different roles in their hosts’ GIT tract. The Testudinimycetales have more limited CAZyme repertoire and a lower temperature range and optima (70, 71), compared to the Neocallimastigales, yet appear to be a predominant and indispensable member of the fungal community in tortoises (family Testudinidae). Their role could be unrelated to food digestion, such as preventing colonization by pathogens via secondary metabolites production. On the other hand, members of the order Neocallimastigales, with the more robust CAZyome, appear to be minor components of the turtle GIT community. As such, their well-known role in the breakdown of plant biomass could be minor or dispensable and could easily by supplanted by bacterial members of the community with broader diversity and temperature optima. Detected sequences of Neocallimastigales in turtles could also be transient components, brought in by the diet of the turtles. Whether Neocallimastigales are a minor but fixed member of the turtle GIT remains to be elucidated. Isolation efforts from fecal samples of turtles outside the family *Testudinidae* are underway.

The chytrid community in Testudines was abundant and diverse, representing 15 orders (Table S2, Figure 3). Chytrid fungi exhibited a lower relative abundance in tortoises (family *Testudinidae*) compared to all other turtle families (Figures 1, 3). Such results could be expected, as the majority of Chytridiomycota families are encountered in aquatic or terrestrial environments, either as saprobic on plants, associated with pollen or diatoms, or pathogenic to plants or algae (78, 100), and hence would have a higher chance of being ingested by water-bound turtles during their feeding.

In addition to providing a detailed description of the fungal community in Testudines, our study expands on the fungal tree of life by identifying additional representatives of multiple recently proposed (77) yet-uncultured phyla (*Candidatus* “Sedimentomastigomycota”, *Candidatus* “Cantoromastigomycota”, *Candidatus* “Tartumycota”, and *Candidatus* “Algovoracomycota”), as well as a few new putative orders, and classes in Chytridiomycota, and Rozellomycota, respectively. It is well established that only a fraction of fungal diversity in nature has been cultured, and that members of the yet-uncultured fungal majority represent vast undescribed biodiversity (101). Indeed, recent culture-independent studies (76, 77, 79, 101, 102) have identified multiple novel lineages in a wide range of habitats, and dedicated curation efforts have allowed placement of such lineages in their appropriate putative taxonomic rank and position in the fungal tree of life (77, 79, 102). In this study, many of the uncultured fungal lineages we identified have been found in a recent meta-analysis of other habitats, mostly soil and sediment (77) (Figures 2, S3–S6). Our effort thus adds additional sequence representatives and new habitats to these cryptic unknown groups in the fungal tree of life. However, we acknowledge that these high-rank taxa appear to be of limited occurrence in our dataset, and when encountered they represented a minor fraction with a few exceptions. For example, while representatives of *Candidatus* “Sedimentomastigomycota” and *Candidatus* “Algovoracomycota” were encountered in 9, and 20 samples, respectively, their relative abundances exceeded 1% in only 1, and 5 samples, respectively, of the 71 samples studied here (Table S2). Similarly, abundances of the new putative orders, and classes in Chytridiomycota, and Rozellomycota, respectively (Figure S4, S6), exceeded 1% in 4 of the 71 samples studied here (Table S2). Therefore, it is uncertain whether high-rank uncultured taxa represent native members of the community in Testudines, are transient in the GIT, or a result of the accidental inclusion of soil in the fecal collection or cloacal swab.

Within animal microbiomes, a fraction of the community is made up of resident members of the community (autochthonous), while the rest represents transient ingested members (allochthonous) from the outside environment (103). The former fraction is expected to be part of the core microbiome of all individuals examined, and exhibit a high level of deterministic selection. Based on our analysis, we argue that members of the Neocallimastigomycota appear to be the only lineage that could convincingly be regarded as host-specific resident members of the community. They were identified in almost all samples (69/71) (Figure 1, Table S2), exhibited strong deterministic patterns of selection (Figure 4), and clearly drove structural differences in mycobiome communities (Figure 5B, S2). Within Neocallimastigomycota, genera in the order Testudinimycetales and *Astrotestudinimyces* genus incerta sedis drove the community separation in tortoises (family *Testudinidae*) (Figure 5B). These Neocallimastigomycota genera represent the oldest lineages in the phylum with evolutionary times ranging from 100–124 Mya (26, 71).

The subgroup Cryptodira harboring the family *Testudinidae* evolved 161–200.2 Mya (104), with the family *Testudinidae* splitting around 51–79 Mya (105). These evolutionary times could argue for a pattern of phylosymbiosis. The Neocallimastigomycota community in other turtle families (order Neocallimastigales), on the other hand, could represent a post-evolutionary environmental filtering process and host selection for taxa from the environment in an evolutionary time-agnostic manner, as genera in Neocallimastigales represent more recent evolutionary times (20–67 Mya) (24), relative to the family *Chelidae* (evolved 101–111 Mya) and the other turtle families (*Emydidae*, *Geoemydidae*, *Kinosternidae*, and *Chelydridae*) that evolved during the late Cretaceous to Early Paleocene (105).

In contrast to Neocallimastigomycota, all other fungal lineages exhibited more stochastic distribution patterns across various hosts, suggesting they represent transient members of the community that were brought about by incidental ingestion. However, we note that, while stochastic in acquisition, the ingested fungal community could further change while passing through the Testudines gut environment, especially given the relatively long residence time (days) (Table S1). For example, the passage through the acidic stomach could result in differential selection of certain taxa that could withstand low pH. Of note, is the high relative abundance of *Candidatus* “Algovoracomycota”, the two uncultured classes of Rozellomycota GS06 and GS137, and the novel orders in Mesochytriomycetes in some samples (Table S2). Members of *Candidatus* “Algovoracomycota” (representatives of the NC_ChyL-1 (also referred to as group GS13)), and members of the Mesochytriomycetes class in Chytridiomycota are thought to be parasites of algae (78, 79), while members of Rozellomycota are known to be obligate intracellular parasites of lower fungi, Oomycetes and algae (78). Parasitism could enable these fungi to be protected inside their hosts while passing through the acidic stomach.

Moreover, post ingestion, some members could not only survive, but possibly thrive and propagate in this new habitat, aided by the change in conditions and relaxation of competition. Two examples from our dataset are noted: the near complete abundance (98.5% of total fungal community) of a Saccharomycetales OTU in one sample (99% similar to *Candida*), and of a Mucorales OTU (83.1% of the total fungal community, 99% similar to *Mucor*) in another. Both genera are ubiquitous facultative anaerobic genera capable of sugar fermentation (106–108), and so would thrive in the anaerobic gut environment.

In conclusion, the current study provides a comprehensive overview of the fungal community in a hitherto understudied reptilian order (Testudines) revealing multiple insights regarding its patterns and determinants. Specifically, it expands the occurrence of the anaerobic gut fungi (Neocallimastigomycota) in non-herbivorous host families, identifies distinct patterns of Neocallimastigomycota community composition in the host family *Testudinidae*, and demonstrates the prevalent role for deterministic processes in shaping the fungal community in family *Testudinidae* as opposed to primarily stochastic processes governing selection in other host families. This study finally broadens the fungal tree of life by identifying multiple novel high-rank yet-uncultured fungal lineages.

## Acknowledgments

This work has been funded by NSF grant 2029478 to MSE and NY, National Science Foundation grants EF-2125065 and CAREER 2236580 to DMW, and NIH grant GM152333 to MSE. Fieldwork in the Philippines was supported by the National Science Foundation DEB 1657648 and IOS 1353683 to CDS and conducted under the Memorandum of Agreement with the Protected Areas and Wildlife Bureau of the Philippines (2015–20), and Gratuitous Permit to Collect #273 (renewal). We thank Mr. Ty Young and Ms. Alexandra Morris, Joseph Spatafora, Kerry McPhail, Jason Stajich, Dale McGinnity at the Nashville Zoo, and Michael Ogle and Steven Nelson at the Knoxville Zoo for assistance with sampling. CDS thanks the Biodiversity Management Bureau (BMB) of the Philippine Department of Environment and Natural Resources (DENR) for help facilitating collecting and export permits, and provincial and municipal authorities in Luzon for facilitating research in regional study sites.

